# Disruption of nasal bacteria enhances protective immune responses to influenza A virus and SARS-CoV-2 infection in mice

**DOI:** 10.1101/2020.12.25.424300

**Authors:** Minami Nagai, Miyu Moriyama, Takeshi Ichinohe

## Abstract

Gut microbiota plays a critical role in the induction of adaptive immune responses to influenza virus infection. However, the role of nasal bacteria in the induction of the virus-specific adaptive immunity is less clear. Here we demonstrate that while intranasal administration of influenza virus hemagglutinin vaccine alone was insufficient to induce the vaccine-specific antibody responses, disruption of nasal bacteria by lysozyme or addition of culturable oral bacteria from a healthy human volunteer rescued inability of the nasal bacteria to generate antibody responses to intranasally administered the split-virus vaccine. Myd88-depdnent signaling in the hematopoietic compartment was required for adjuvant activity of intranasally administered oral bacteria. In addition, we found that the oral bacteria-combined intranasal vaccine induced protective antibody response to influenza virus and SARS-CoV-2 infection. Our findings here have identified a previously unappreciated role for nasal bacteria in the induction of the virus-specific adaptive immune responses.

## Introduction

Respiratory infectious diseases such as influenza and coronavirus disease 2019 (COVID-19) cause substantial morbidity and mortality. Influenza A virus is responsible for annual epidemics that cause severe morbidity and mortality involving 3 to 5 million people worldwide. In addition, the constant pandemic potential of newly emerging viruses remains a serious threat to public health, the economy and society as illustrated by the recent COVID-19 global pandemic. Therefore, there is an urgent need to develop effective vaccines against not only seasonal influenza viruses but also against severe acute respiratory syndrome coronavirus 2 (SARS-CoV-2).

Since it is difficult to predict which strain of influenza virus or coronavirus cause a pandemic, it is advantageous to produce vaccines that induce cross-protective immunity against variants of the particular virus strain. Mucosal immunity induced by natural infection of influenza virus is more effective and cross-protective against heterologous virus infection than systemic immunity induced by parenteral vaccines (S. Tamura & Kurata, 2004). It is believed that the virus-specific IgA in upper respiratory tract is more cross-protective against heterologous influenza viruses compared with the virus-specific IgG in the serum due to its dimeric or tetrameric forms (higher avidity) and location (Liew, Russell, Appleyard, Brand, & Beale, 1984; Suzuki et al., 2015). Indeed, polymeric immunoglobulin receptor-knockout mice failed to secrete nasal IgA and protect against heterologous virus challenge (Asahi et al., 2002). Therefore, induction of the virus-specific secretory IgA in the upper respiratory tract by intranasal vaccination has a great advantage in conferring protection against an unpredictable pandemic of viral pathogens such as the swine-origin H1N1 and avian-origin H7N9 influenza A viruses, or zoonotic origin of SARS-CoV-2 (Gao et al., 2013; Neumann, Noda, & Kawaoka, 2009). In the effort to develop effective intranasal vaccines, several adjuvants such as cholera toxin (Watanabe et al., 2002), synthetic double-stranded RNA poly(I:C) (Ichinohe et al., 2005), synthetic toll-like receptor 4 agonist (Spinner et al., 2015), zymosan (Ainai et al., 2010), flagellin (Skountzou et al., 2010), immune stimulating complexes (ISCOMs) (Sjolander et al., 2001), or type-I interferons (Bracci et al., 2005) have been developed to enhance the vaccine-specific nasal IgA response. While upper respiratory tract contains commensal bacteria (Bassis et al., 2015; Clark, 2020), intranasal administration of split vaccine alone was insufficient to induce the vaccine-specific nasal IgA response (Ichinohe et al., 2005; Jangra et al., 2020), suggesting that the amounts of commensal bacteria in upper respiratory tract are insufficient to stimulate the vaccine-specific nasal IgA response.

A recent study has demonstrated that nasal mucosa-derived *Staphylococcus epidermidis*, one of the most abundant colonizers of healthy human skin and mucosal surface, suppressed influenza virus replication by stimulating IFN-λ production (Kim et al., 2019). In addition, influenza virus-infected mice lacking both toll-like receptor 7 (TLR7) and mitochondrial antiviral signaling (MAVS) had elevated nasal bacterial burdens, which resulted in death from pneumonia caused by secondary bacterial infections (Pillai et al., 2016). In contrast to the role of nasal bacteria in innate antiviral resistance to influenza virus infection or severity of the disease (Kim et al., 2019; Pillai et al., 2016), it remains unclear whether nasal bacteria critically regulates the generation of influenza virus-specific adaptive immune responses after infection or intranasal vaccination. Here, we show that depletion of nasal bacteria by intranasal administration of antibiotics enhanced the virus-specific nasal IgA and serum IgG response following influenza virus infection. In addition, we found that lysozyme-induced disruption of nasal bacteria or culturable oral bacteria from a healthy volunteer significantly enhanced the vaccine-specific nasal IgA and serum IgG responses. Myd88-depdnent signaling in the hematopoietic compartment was required for adjuvant activity of intranasally administered oral bacteria. Our findings here have identified a previously unappreciated role for nasal bacteria in the induction of the virus-specific adaptive immune responses.

## Results

### Depletion of nasal bacteria enhanced antibodies response to influenza virus infection

Gut commensal microbiota play a key role in innate and adaptive immune defense against influenza virus infection (Abt et al., 2012; Bradley et al., 2019; Ichinohe et al., 2011; Rosshart et al., 2017; Steed et al., 2017; Stefan, Kim, Iwasaki, & Kasper, 2020). However, the role of oral or nasal bacteria in the induction of mucosal immune responses following influenza virus infection remains unknown. To assess the effects of oral or nasal bacteria in the induction of mucosal immune responses to influenza virus infection, we treated mice intranasally with an antibiotic cocktail for five consecutive days before influenza virus infection. This treatment resulted in significant reduction in the numbers of culturable oral and nasal bacteria (**Supplementary Fig. 1**).

Antibiotic-treated mice were then infected intranasally with a mouse-adapted influenza A virus strain A/Puerto Rico/8/1934 (PR8). Surprisingly, influenza virus-specific nasal IgA and serum IgG levels were significantly elevated in the antibiotic-treated group (**Fig. 1**). This led us to consider the possibility that depletion of commensal bacteria in upper respiratory tract enhances influenza virus replication, resulting in enhancement of the virus-specific antibody responses. However, depletion of commensal bacteria in upper respiratory tract significantly reduced influenza virus replication at 2 days post infection (**Supplementary Fig. 2A**). This is consistent with a previous report showing that antibiotic treatment significantly reduce influenza virus replication at early time point (Gopinath et al., 2018). In addition, the viral replication in upper respiratory tract became comparable between antibiotics-treated and control groups at 3 and 5 days post infection (**Supplementary Fig. 2B, C**). These data indicated that the levels of influenza virus replication in upper respiratory tract is unlikely to account for increased the virus-specific antibody responses in antibiotic-treated animals.

**Figure 1.**
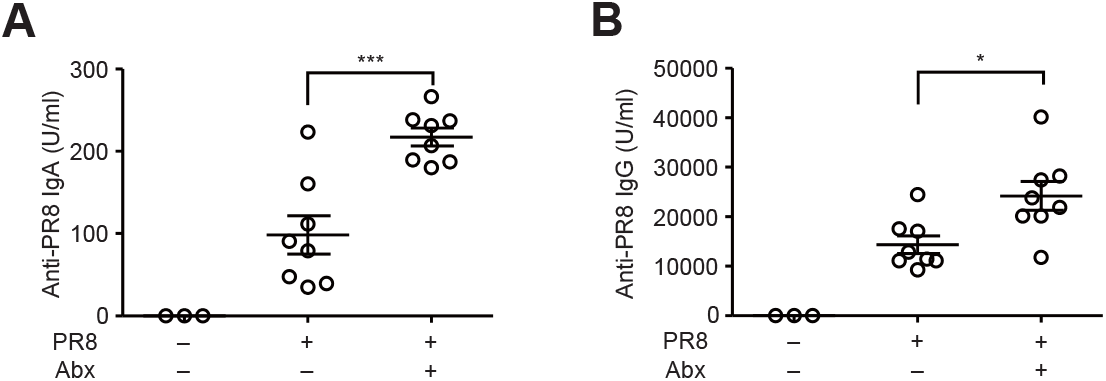
Disruption of nasal bacteria enhances the virus-specific antibody responses following influenza virus infection. (**A and B**) Mice were inoculated intranasally with an antibiotic cocktail (Abx) for 5 consecutive days. Two days later, mice were intranasally infected with 1,000 pfu of A/PR8 virus. The nasal wash and serum were collected at 4 weeks p.i., and the virus-specific nasal IgA and serum IgG titers were determined by ELISA. Open circles indicate values for individual mice. The data are from three independent experiments (mean ± SEM). \**P* < 0.05 and \*\*\**P* < 0.001; (one-way ANOVA and Tukey’s test).

### Lysozyme-induced disruption of nasal bacteria enhances antibody responses induced by intranasal vaccination

Thus, we next examined the possibility that antibiotic-induced disruption of nasal bacteria releases pathogen-associated molecular patterns, which may act as adjuvants to enhance the virus-specific antibody responses. To assess the possibility that disruption of nasal bacteria acts as adjuvant for intranasal influenza vaccine, we immunized mice intranasally with influenza virus hemagglutinin (HA) protein and lysozyme to disrupt nasal bacteria. We used poly(I:C) adjuvant as a positive control (Ichinohe et al., 2005). Strikingly, we found that intranasal immunization with HA and lysozyme significantly enhanced the HA-specific nasal IgA and serum IgG responses (**Fig. 2**). While upper respiratory tract contains commensal bacteria (Bassis et al., 2015; Clark, 2020), intranasal administration of hemagglutinin (HA) vaccine alone was insufficient to induce the HA-specific antibody responses (**Fig. 2**). Taken together, these results suggest that disruption of nasal bacteria by intranasal administration of antibiotics or lysozyme acts as adjuvant for intranasal influenza vaccine.

**Figure 2.**
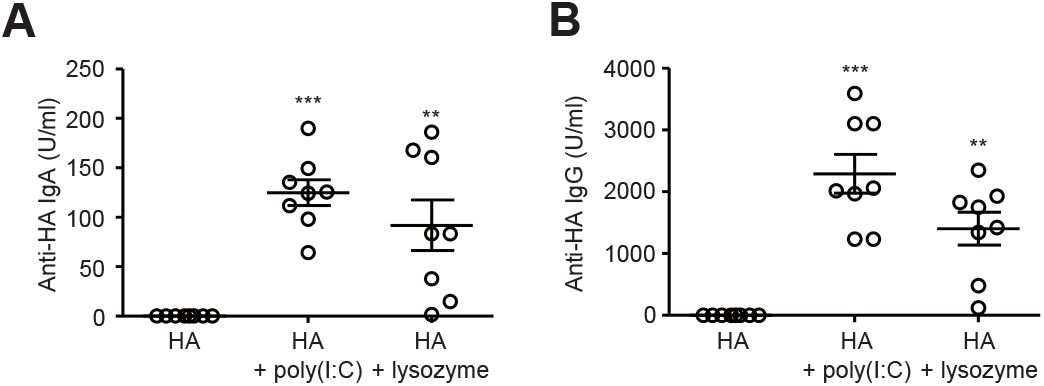
Disruption of nasal bacteria induces the HA-specific antibody responses after intranasal vaccination. (**A and B**) Mice were immunized intranasally with quadrivalent HA vaccine with or without poly(I:C) or lysozyme twice in a 3-week interval. Two weeks later, the nasal wash and serum were collected and the HA-specific nasal IgA and serum IgG titers were determined by ELISA. Open circles indicate values for individual mice. The data are from three independent experiments (mean ± SEM). \*\**P* < 0.01 and \*\*\**P* < 0.001; (one-way ANOVA and Tukey’s test).

### Oral bacteria act as adjuvant for intranasal vaccine

While upper respiratory tract contains commensal bacteria (Bassis et al., 2015; Clark, 2020), we found that relative amounts of 16S rRNA and culturable bacteria in nasal mucosal surface were significantly lower than that in the oral cavity (**Supplementary Fig. 3**). Thus, we next examine whether oral bacteria act as adjuvant for intranasal vaccine. Intranasal vaccination with HA and culturable oral bacteria from mice or a healthy volunteer significantly enhanced the HA-specific nasal IgA and serum IgG responses (**Fig. 3A, B**). In addition, the oral bacteria from a healthy volunteer stimulated the HA-specific nasal IgA and serum IgG responses in a dose-dependent manner (**Fig. 3C, D**). Next, we compared the ability of isolated bacterial strains from oral wash sample of a healthy volunteer to stimulate the HA-specific antibody responses. To this end, we immunized mice intranasally with HA and *streptococcus salivarius* (*S. salivarius*), *streptococcus parasanguinis* (*S. parasanguinis*), or *streptococcus infantis* (*S. infantis*). Mice immunized with HA and each isolated bacterial strain induced comparable levels of the HA-specific nasal IgA and serum IgG responses (**Fig. 4**), suggesting that adjuvant activity of the oral bacteria is unlikely to account for strain specific.

**Figure 3.**
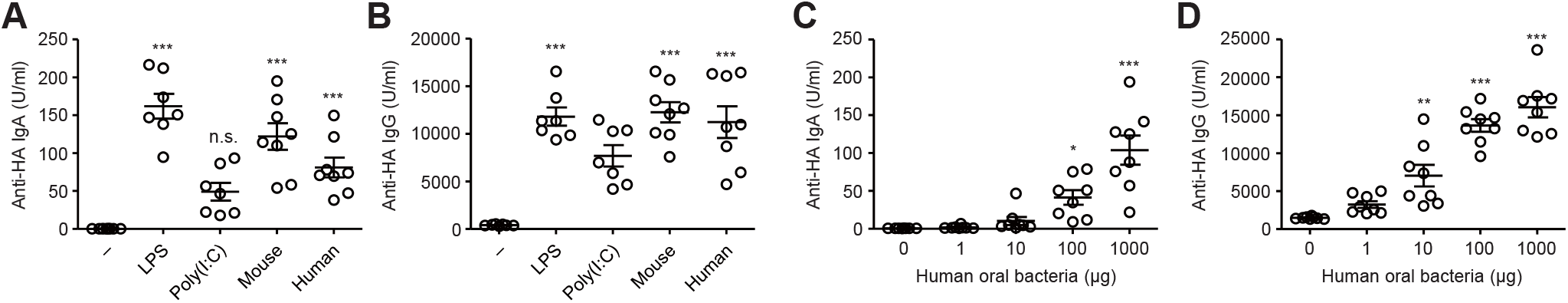
Oral bacteria acts as adjuvant for intranasal vaccine. (**A and B**) Mice were immunized intranasally with quadrivalent HA vaccine with or without LPS, poly(I:C), or culturable oral bacteria from mice or a healthy volunteer twice in a 3-week interval. Two weeks later, the nasal wash and serum were collected and the HA-specific nasal IgA and serum IgG titers were determined by ELISA. (**C and D**) Mice were immunized intranasally with quadrivalent HA vaccine with or without indicated amounts of oral bacteria from a healthy volunteer twice in a 3-week interval. Two weeks later, the nasal wash and serum were collected and the HA-specific nasal IgA and serum IgG titers were determined by ELISA. Open circles indicate values for individual mice. The data are from two independent experiments (mean ± SEM). \**P* < 0.05, \*\**P* < 0.01 and \*\*\**P* < 0.001; (one-way ANOVA and Tukey’s test).

**Figure 4.**
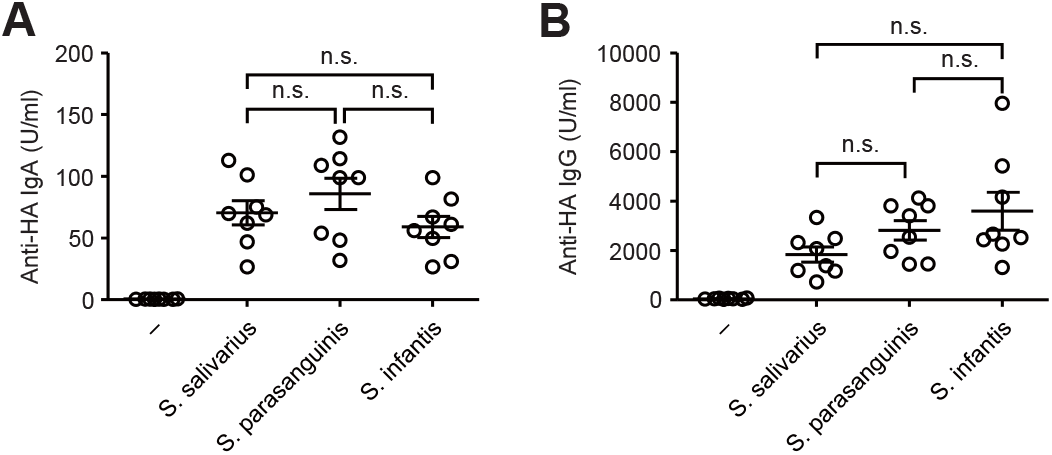
Adjuvant activity of *S. salivarius*, *S. parasanguinis*, and *S. infantis* for intranasal vaccine. (**A and B**) Mice were immunized intranasally with quadrivalent HA vaccine with or without *S. salivarius*, *S. parasanguinis*, or *S. infantis* twice in a 3-week interval. Two weeks later, the nasal wash and serum were collected and the HA-specific nasal IgA and serum IgG titers were determined by ELISA. Open circles indicate values for individual mice. The data are from two independent experiments (mean ± SEM). \**P* < 0.05, \*\**P* < 0.01 and \*\*\**P* < 0.001; (one-way ANOVA and Tukey’s test).

### Myd88-depdnent signaling in the hematopoietic compartment is required for adjuvant activity of intranasally administered oral bacteria

Next, we wished to determine the innate immune signaling through pattern-recognition receptors required for adjuvant activity of the oral bacteria. To this end, we immunized WT and MyD88-deficient mice intranasally with HA and culturable oral bacteria from a healthy volunteer and measured the HA-specific nasal IgA and serum IgG responses. The HA-specific nasal IgA and serum IgG responses were found to be completely dependent on MyD88 (**Fig. 5A, B**). In addition, lysozyme-induced disruption of nasal bacteria stimulated the HA-specific nasal IgA and serum IgG responses in a MyD88-dependent manner (**Fig. 5C, D**). To determine the cellular compartment responsible for adjuvant activity of oral bacteria, we generated bone marrow (BM) chimeric mice in which only the hematopoietic (WT→MyD88^−/−^) or the stromal cells (MyD88^−/−^→WT) expressed MyD88. After intranasal vaccination with HA and oral bacteria, the HA-specific nasal IgA and serum IgG responses were significantly reduced in MyD88^−/−^→WT BM chimeric mice compared to WT→MyD88^−/−^ BM chimeric mice (**Fig. 6**). These data indicate that MyD88-dependent signaling in the hematopoietic, but not stromal, compartment is required for adjuvant activity of intranasally administered oral bacteria.

**Figure 5.**
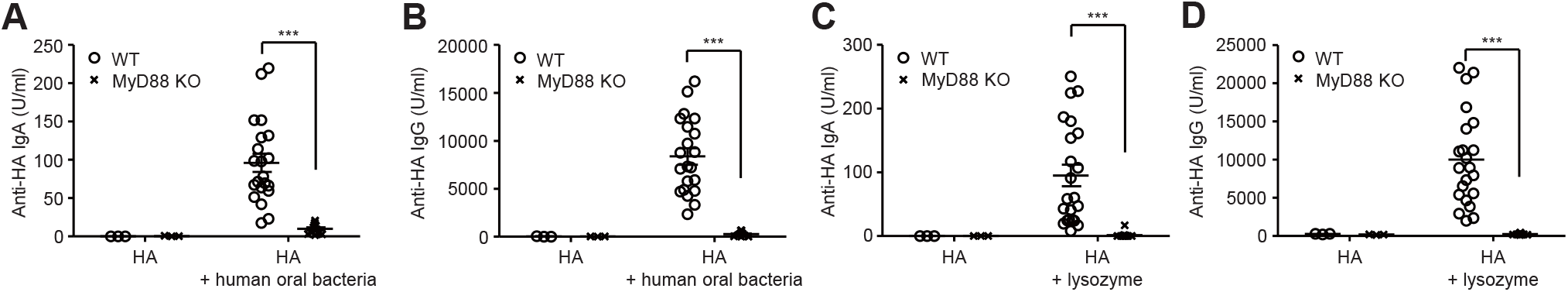
Oral bacteria acts as adjuvant for intranasal vaccine. (**A-D**) WT and MyD88-deficient mice were immunized intranasally with quadrivalent HA vaccine with or without culturable oral bacteria from a healthy volunteer (**A and B**) or lysozyme (**C and D**) twice in a 3-week interval. Two weeks later, the nasal wash and serum were collected and the HA-specific nasal IgA and serum IgG titers were determined by ELISA. Open circles indicate values for individual mice. The data are from two independent experiments (mean ± SEM). \*\*\**P* < 0.001; (one-way ANOVA and Tukey’s test).

**Figure 6.**
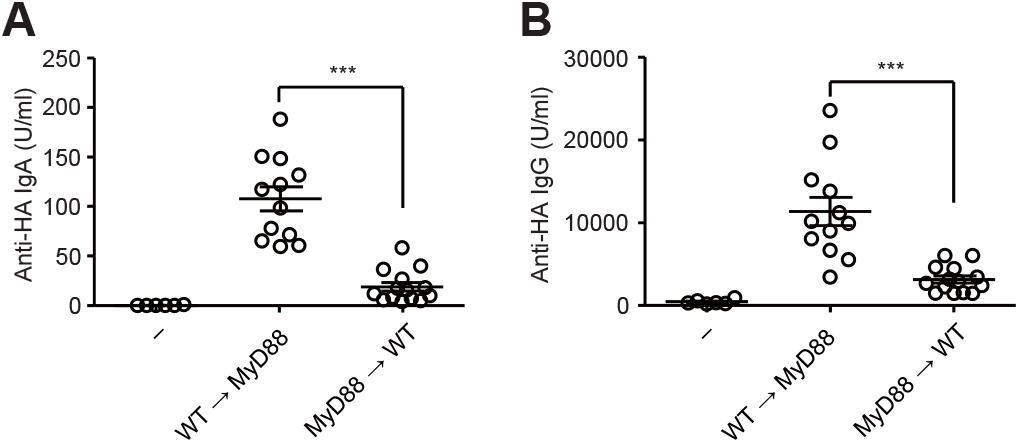
Oral bacteria acts as adjuvant for intranasal vaccine. (**A and B**) WT→MyD88 KO and MyD88 KO→WT BM chimeric mice were immunized intranasally with quadrivalent HA vaccine with or without culturable oral bacteria from a healthy volunteer twice in a 3-week interval. Two weeks later, the nasal wash and serum were collected and the HA-specific nasal IgA and serum IgG titers were determined by ELISA. Open circles indicate values for individual mice. The data are from two independent experiments (mean ± SEM). \*\*\**P* < 0.001; (one-way ANOVA and Tukey’s test).

### Oral bacteria-combined intranasal vaccine protects from influenza virus and SARS-CoV-2 infection

Finally, we examined protective effects of intranasal vaccination with oral bacteria-adjuvanted vaccine against influenza virus and SARS-CoV-2 infection. To this end, we immunized mice intranasally with quadrivalent influenza HA vaccine containing A/California/7/2009 HA together with culturable oral bacteria or lysozyme. Two weeks after the second vaccination, we challenged vaccinated mice intranasally with a heterologous A/Narita/1/2009 (pdm09) strain (**Fig. 7**). Mice immunized with HA vaccine adjuvanted with oral bacteria or lysozyme significantly reduced the virus titer compared to control mice immunized with the HA vaccine alone (**Fig. 7**). We next assessed protective effects of intranasal vaccination with oral bacteria-adjuvanted SARS-CoV-2 spike protein against SARS-CoV-2 infection in Syrian hamsters. To this end, we immunized hamsters intranasally with a recombinant SARS-CoV-2 spike protein and culturable oral bacteria from a healthy volunteer. We immunized hamsters subcutaneously with the spike protein alone as a control. We Both the spike- and the virus-specific serum IgG levels were significantly elevated in immunized hamsters (**Fig. 8A, B**). In addition, immunized hamsters significantly reduced the virus titer compared to naïve animals following high-dose (2×10^6^ pfu of SARS-CoV-2) challenge (**Fig. 8C**). These data collectively indicated that disruption of nasal bacteria or intranasal administration of oral bacteria compensate inability of nasal bacteria to generate protective adaptive immunity to intranasally administered split vaccines.

**Figure 7.**
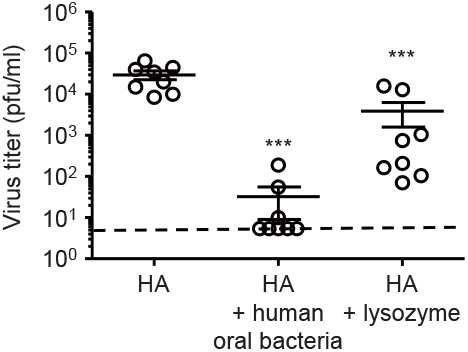
Protective effects of oral bacteria-adjuvanted intranasal vaccine against influenza virus infection. Mice were immunized intranasally with quadrivalent HA vaccine with or without culturable oral bacteria from a healthy volunteer or lysozyme twice in a 3-week interval. Two weeks after the last vaccination, mice were challenged with 1,000 pfu of A/PR8 virus. The nasal wash of influenza virus-infected mice was collected at 3 days post infection, and viral titers were determined by plaque assay. Open circles indicate values for individual mice. The dashed line indicates the limit of virus detection. The data are from two independent experiments (mean ± SEM). \*\*\**P* < 0.001; (one-way ANOVA and Tukey’s test).

**Figure 8.**
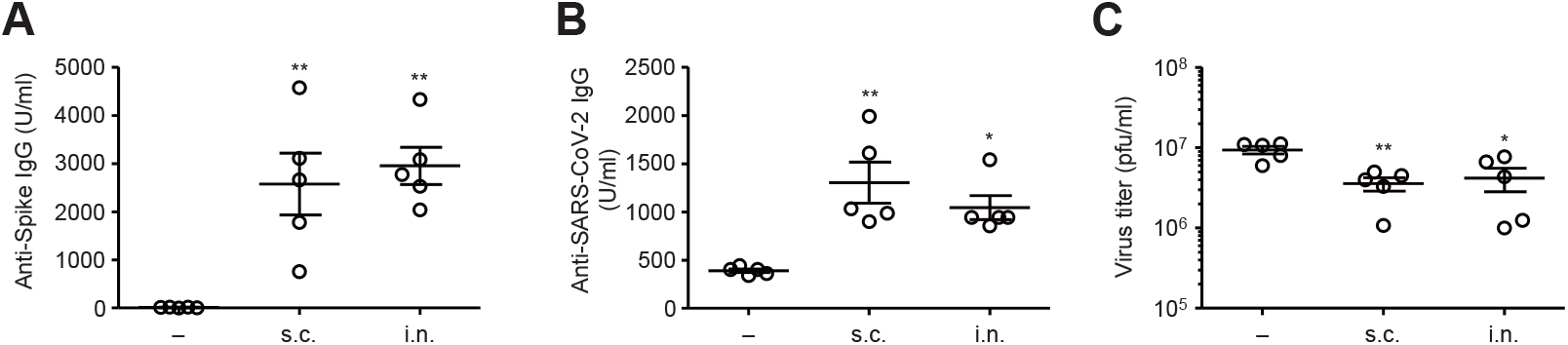
Protective effects of oral bacteria-adjuvanted intranasal vaccine against SARS-CoV-2 infection. (**A-C**) Hamsters were immunized subcutaneously or intranasally with the spike protein of SARS-CoV-2 with or without culturable oral bacteria from a healthy volunteer twice in a 3-week interval. Two weeks after the last vaccination, hamsters were challenged with 2×10^6^ pfu of SARS-CoV-2. (**A and B**) Serum were collected at 3 days post infection. The spike protein- (**A**) or SARS-CoV-2- (**B**) specific serum IgG antibody titers were determined by ELISA. (**C**) The lung wash of SARS-CoV-2-infected hamsters was collected at 3 days post infection, and viral titers were determined by plaque assay. Open circles indicate values for individual hamsters. The data are from two independent experiments (mean ± SEM). \**P* < 0.05 and \*\**P* < 0.01; (one-way ANOVA and Tukey’s test).

## Discussion

The innate immune system, the first line of defense against pathogens, utilizes pattern recognition receptors (PRRs) to detect pathogen-associated molecular patterns (PAMPs). The recognition of influenza virus by PRRs plays a key role not only in limiting virus replication at early stages of infection, but also in initiating the virus-specific adaptive immune responses. In addition, previous studies have demonstrated that gut commensal microbiota play a key role in innate and adaptive immune defense against influenza virus infection (Abt et al., 2012; Bradley et al., 2019; Ichinohe et al., 2011; Rosshart et al., 2017; Steed et al., 2017; Stefan et al., 2020). Further, recent studies have indicated the roles of nasal bacteria in innate antiviral resistance to influenza virus infection or severity of the diseases (Kim et al., 2019; Pillai et al., 2016). However, it remains unclear whether nasal bacteria critically regulates the generation of influenza virus-specific adaptive immune responses after influenza virus infection. In this study, we demonstrated that depletion of commensal bacteria in upper respiratory tract by intranasal administration of antibiotics enhanced the virus-specific antibodies response following influenza virus infection. Surprisingly, depletion of nasal bacteria by intranasal administration of antibiotics before influenza virus infection significantly reduced the virus titer at 2 days post infection. This is consistent with a previous report showing that antibiotic treatment significantly reduce influenza virus replication at 6 hours post infection (Gopinath et al., 2018). Intranasal application of antibiotics suppressed influenza virus replication through at least two possible mechanisms. First, intranasal administration of antibiotics enhances host resistance to influenza virus infection in a microbiota-independent manner (Gopinath et al., 2018). Second, disruption of nasal bacteria by intranasal antibiotic treatment may release PAMPs from the antibiotic-killed bacteria, which stimulate innate antiviral immune responses to suppress influenza virus replication (Matsuo et al., 2000). After 3 and 5 days post infection, the viral replication in upper respiratory tract became comparable between antibiotic-treated and control groups, indicating that the levels of influenza virus replication in upper respiratory tract is unlikely to account for increased levels of the virus-specific antibodies response in antibiotic-treated mice.

Since the primary targets of influenza virus are the nasal epithelial cells in upper respiratory tract, it is beneficial to induce the virus-specific nasal IgA antibody at the nasal mucosal epithelium. However, intranasal vaccination with split-virus vaccine alone is often insufficient to elicit proper immune responses at the upper respiratory tract. Therefore, adjuvants are required for a given vaccine to induce the vaccine-specific nasal IgA response. In developing intranasal vaccines, cholera toxin (CT) and *Escherichia coli* heat-labile toxin (LT) have been used as adjuvant to enhance nasal immune response (S. I. Tamura & Kurata, 2000). Although CT and LT are effective adjuvants to enhance mucosal immune responses including secretory IgA responses, they have some side effects in humans, including Bell’s palsy and nasal discharge (Mutsch et al., 2004). Therefore, several adjuvants that are as effective as CT or LT and are also safe for human use have been developed for clinical application with intranasal influenza vaccine (Ainai et al., 2010; Bracci et al., 2005; Ichinohe et al., 2005; Sjolander et al., 2001; Skountzou et al., 2010; Spinner et al., 2015). In this study, we show that intranasal vaccination with influenza virus HA vaccine and culturable oral bacteria from a healthy human volunteer induced significant levels of the vaccine-specific nasal IgA and serum IgG responses in a dose-dependent manner. All commensal bacterial strains tested, including *S. salivarius*, *S. parasanguinis*, or *S. infantis*, induced comparable levels of the HA-specific nasal IgA and serum IgG responses, suggesting that adjuvant activity of the oral bacteria is unlikely to account for strain specific. In addition to culturable oral bacteria from a healthy human volunteer, we demonstrated that disruption of nasal bacteria by lysozyme induced significant levels of the vaccine-specific antibodies response. Although relative amounts of nasal bacteria were significantly lower than that in the oral cavity, disruption of nasal bacteria by lysozyme could rescue the inability of nasal bacteria to generate the vaccine-specific antibodies response. In mice, nasal commensal microbiota are predominantly composed of gram-positive bacteria including *Lactobacillus spp*., *Bacillus spp*., *Staphylococcus spp*., and *Streptococcus spp*. (Ichinohe et al., 2011). In addition, *Lactobacillus spp*. were found to contain higher amounts of double-stranded RNA than the pathogenic bacteria (Kawashima et al., 2013). Since activation of TLRs by different PAMPs such as poly(I:C) and zymosan synergistically enhanced the nasal IgA response to intranasally administered influenza virus HA vaccine (Ainai et al., 2010), disruption of nasal bacteria could stimulate different TLRs to enhance the vaccine-specific antibodies response. Most TLRs signal though the adaptor protein MyD88 (Kawai & Akira, 2010; Medzhitov, 2001). Although nasal epithelial cells express various TLRs (Tengroth et al., 2014; van Tongeren et al., 2015), deficiency of MyD88 in stromal compartment did not significantly affect the levels of nasal IgA and serum IgG responses following intranasal vaccination with influenza virus HA and culturable oral bacteria. Instead, MyD88-dependent signaling in the hematopoietic cells were required for adjuvant activity of intranasally administered oral bacteria. These data are consistent with previous studies showing that both TLR-induced dendritic cell maturation and B-cell activation are required for optimal antibody responses to T-dependent antigens (Iwasaki & Medzhitov, 2015; Pasare & Medzhitov, 2005).

In summary, our study demonstrated the effects of commensal microbiota in upper respiratory tract in the induction of the virus-specific adaptive immune responses after influenza virus infection or intranasal vaccination. Our data indicated that disruption of nasal bacteria by lysozyme or supplementation of oral bacteria from a healthy volunteer enhanced nasal IgA and serum IgG antibodies response to intranasally administered influenza virus HA or SARS-CoV-2 S proteins. Although the vaccinated animals significantly reduced the virus titer compared to unadjuvanted group or naïve animals following high-dose of influenza virus or SARS-CoV-2 challenge, further studies are needed to establish the safety and efficacy of this vaccination method in an additional animal model such as nonhuman primate.

## Materials and methods

### Mice

Age- and sex-matched Balb/c mice obtained from Japan SLC, Inc. were used as WT controls. MyD88-deficient Balb/c mice were a gift from T. Taniguchi. All animal experiments were performed in accordance with the University of Tokyo’s Regulations for Animal Care and Use, which were approved by the Animal Experiment Committee of the Institute of Medical Science, the University of Tokyo (approval number PA17– 69).

### Cells

Madin-Darby canine kidney (MDCK) cells were grown in Eagle’s minimal essential medium (E-MEM; Nacalai Tesque) supplemented with 10% fetal bovine serum (FBS), penicillin (100 U/ml), and streptomycin (100 μg/ml). VeroE6 cells stably expressing transmembrane protease serine 2 (VeroE6/TMPRSS2; JCRB Cell Bank 1819) were maintained in Dulbecco’s modified Eagle’s medium (DMEM) low glucose (Cat#08456-65; Nacalai Tesque) supplemented with 10% FBS, penicillin (100 U/ml), streptomycin (100 μg/ml), and G418 (1mg/ml) (Matsuyama et al., 2020).

### Depletion of nasal bacteria *in vivo*

The antibiotic cocktail consisted of ampicillin sodium salt (1 g/L), neomycin sulfate (1 g/L), metronidazole (1 g/L), vancomycin hydrochloride (0.5 g/L), gentamicin (10 mg/L), penicillin (100 U/ml), streptomycin (100 U/ml), and amphotericin B (0.25 mg/L) (Moriyama & Ichinohe, 2019). For intranasal treatment, mice were anaesthetized and 5 μl of antibiotic was administered dropwise into each nostril using a pipette tip. All antibiotics with the exception of vancomycin hydrochloride were obtained from Nacalai Tesque. Vancomycin hydrochloride was obtained from Duchefa Biochemie.

### Virus infection

WT A/Puerto Rico/8/34 (A/PR8) and A/Narita/1/09 (pdm09) influenza viruses were grown in allantoic cavities of 10-d-old fertile chicken egg at 35°C for 2 d (Moriyama et al., 2020). Viral titer was quantified by a standard plaque assay using MDCK cells and viral stock was stored at –80°C (Moriyama, Koshiba, & Ichinohe, 2019). For intranasal infection, mice were fully anesthetized by i.p. injection of pentobarbital sodium (Somnopentyl, Kyoritsu Seiyaku Co., Ltd., Tokyo, Japan) and then infected by intranasal application of 30 μl of virus suspension (1,000 pfu of A/PR8 or pdm09 in PBS). This procedure leads to upper and lower respiratory tract infection (Moriyama & Ichinohe, 2019).

SARS-CoV-2 (a gift from Y. Kawaoka) was amplified on VeroE6/TMPRSS2 cells and stored at –80°C until use. The infectious titer was determined by a standard plaque assay using VeroE6/TMPRSS2 cells, as described previously (Imai et al., 2020). For intranasal infection, one-month-old female Syrian hamsters (Japan SLC Inc.) were fully anesthetized by i.p. injection of pentobarbital sodium (Somnopentyl, Kyoritsu Seiyaku Co., Ltd., Tokyo, Japan) and then infected intranasally with 2×10^6^ pfu (in 100 μL) of SARS-CoV-2. All experiments with SARS-CoV-2 were performed in enhanced biosafety level 3 (BSL-3) containment laboratories at the University of Tokyo, in accordance with the institutional biosafety operating procedures.

### Vaccination

For intranasal infection, mice were fully anesthetized by i.p. injection of pentobarbital sodium (Somnopentyl, Kyoritsu Seiyaku Co., Ltd., Tokyo, Japan) and then infected intranasally by dropping 2 μl of PBS containing 1,000 pfu of A/PR8 into the nostril. The quadrivalent inactivated influenza vaccine (split-product virus vaccines, hemagglutinin [HA] vaccine) prepared for the 2015–2016 season and including A/California/7/2009 (H1N1), A/Switzerland/9715293/2013 (H3N2), B/Phuket/3073/2013, and B/Texas/2/2013 were purchased from Kaketsuken (Kumamoto, Japan). Mice were immunized by intranasal administration of the quadrivalent HA vaccine containing 150 ng of each HA with or without 5 μg of lipopolysaccharide (LPS; InvivoGen), 5 μg of poly(I:C) (InvivoGen), 250 μg of lysozyme (Thermo Fisher Scientific), or 1 mg of culturable oral bacteria from a healthy volunteer.

SARS-CoV-2 spike S1+S2 ECD-His recombinant protein was purchased from Sino Biological Inc. (Cat# 40589-V08B1). Hamsters were immunized subcutaneously or intranasally with 1 μg of the recombinant spike protein with or without 1 mg of culturable oral bacteria from a healthy volunteer.

### Clinical specimens

Oral and nasal washes were collected from a healthy volunteer by rinsing the mouth with 50 ml of saline or washing the nasal cavity with 50 ml of saline using a syringe. The research protocol was approved by the Research Ethics Review Committee of the Institute of Medical Science, the University of Tokyo (approval number 2019-42-1121). For preparation of oral bacteria adjuvant, oral wash samples were grown in brain heart infusion broth (BD 237500) at 37°C overnight, washed repeatedly, and resuspended in PBS (200 μg/ml).

### Bacterial recovery and identification

Oral and nasal washes were collected from a healthy volunteer as described above. Aliquots of 100μl of serial 10-fold dilution of the oral and nasal wash were inoculated into brain heart infusion agar plates (BD 252109). After incubation at 37°C overnight under the aerobic conditions, the bacterial colonies were grown in brain heart infusion broth (BD 237500) at 37°C overnight. Bacterial DNA was isolated as described previously (Moriyama & Ichinohe, 2019). A 300-bp portion of the 16S rRNA was amplified by PCR using specific primer pairs of 515F (5’-GTGCCAGCMGCCGCGGTAA-3’) and 806R (5’-GGACTACHVGGGTWTCTAAT-3’), purified (Qiagen), sequenced, and the sequence compared by Blast analysis to known bacterial sequences.

### Bone marrow chimera

Bone marrow chimeras were generated as described (Pang, Ichinohe, & Iwasaki, 2013). WT and MyD88-deficient mice were γ-irradiated with 6 Gy, then were reconstituted with 5 × 10^6^ bone marrow cells of the appropriate genotype via i.v. injection and allowed to recover for 8 weeks before vaccination.

### Measurement of virus titers

For measurement of influenza virus titer, bronchoalveolar (BAL) fluid was collected by washing the trachea and lungs of mice twice by injecting a total of 2 ml PBS containing 0.1% bovine serum albumin (BSA). The virus titer was measured as follows: aliquots of 200 μl of serial 10-fold dilution of the BAL fluid by PBS containing 0.1% BSA were inoculated into MDCK cells in 6-well plates. After 1 hour of incubation, cells were washed with PBS thoroughly and overlaid with 2 ml of agar medium.

For measurement of SARS-CoV-2 titer, BAL fluid was collected by washing the trachea and lungs of hamsters twice by injecting a total of 2 ml DMEM containing 5% FBS. The virus titer was measured as follows: aliquots of 200 μl of serial 10-fold dilution of the BAL fluid by DMEM containing 5% FBS were inoculated into VeroE6/TMPRSS2 cells in 6-well plates. After 1 hour of incubation, cells were washed with PBS thoroughly and overlaid with 2 ml of agar medium. The number of plaques in each well was counted 2 days after inoculation.

### Enzyme-linked immunosorbent assay (ELISA)

Serum and nasal wash were collected from the immunized mice for measurement of the PR8- or HA-specific nasal IgA and serum IgG antibodies. Nasal wash was collected by washing the nasopharynx three times by injecting a total of 1 ml PBS containing 0.1% BSA. The levels of the PR8- or HA-specific nasal IgA and serum IgG antibodies were determined by ELISA as described previously (Moriyama & Ichinohe, 2019). Standards for PR8- or HA-reactive IgA and IgG antibody titration were prepared from the nasal wash or serum of the virus-infected or vaccinated mice, and expressed as the same arbitrary units (160-unit). The antibody titers of unknown specimens were determined from the standard regression curve constructed by two fold serial dilution of the 160-unit standard for each assay.

### Quantification and statistical analysis

Statistical significance was tested by one-way ANOVA followed by Tukey test or unpaired t tests with PRISM software (Version 5; GraphPad software). Data are presented as mean ± SEM. Statistical details can be found directly in the figure legends. P values of less than 0.05 were considered statistically significant.

## ACKNOWLEDGMENTS

We thank Y. Kawaoka (University of Wisconsin and University of Tokyo) for providing SARS-CoV-2 and T. Taniguchi (The University of Tokyo) for MyD88-deficient mice. This work was supported by the Japan Society for the Promotion of Science Grants-in-Aid for Scientific Research (20H03491), the Research Program on Emerging and Re-emerging Infectious Diseases, of the Japan Agency for Medical Research and Development (AMED), the Yakult Bio-Science Foundation, the Hitachi Global Foundation, and the Mitsubishi foundation. M. M. is the Research Fellow of the Japan Society for the Promotion of Science.

