## Supplementary Figures for "Disruption of nasal bacteria enhances protective immune responses to influenza A virus and SARS-CoV-2 infection in mice"

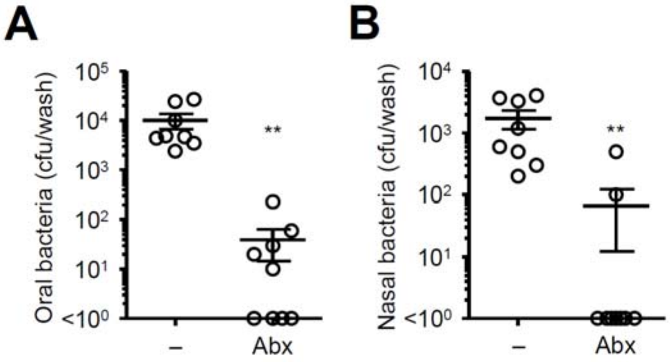


**Figure S1. Effect of intranasal antibiotic treatment on oral and nasal bacterial load.** Mice were inoculated intranasally with an antibiotic cocktail (Abx) for 5 consecutive days. Two days later, tongue (A) and nasal wash (B) were collected by washing the nasopharynx three times by injecting a total of 1 ml of brain heart infusion broth. Bacterial load in the tongue (A) and nasal wash (B) were measured. **P < 0.01; (one-way ANOVA and Tukey's test).

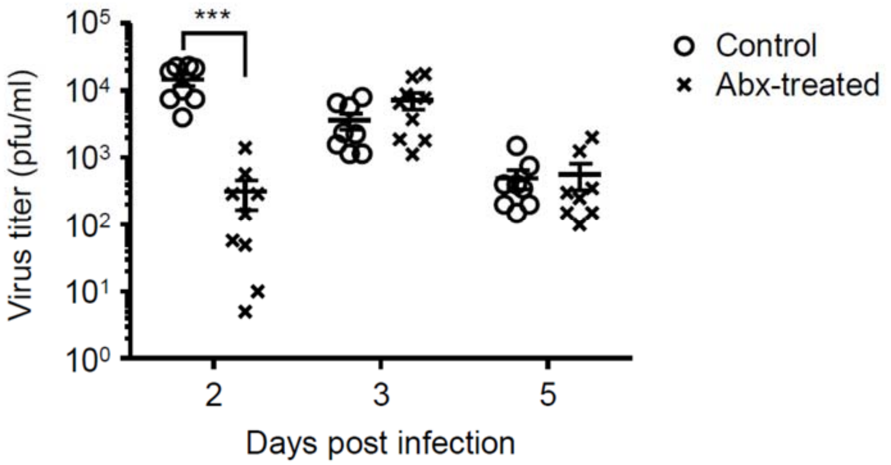


**Figure S2. Effect of intranasal antibiotic treatment on influenza virus replication.** Mice were inoculated intranasally with an antibiotic cocktail (Abx) for 5 consecutive days. Two days later, mice were intranasally infected with 1,000 pfu of A/PR8 virus. The nasal wash was collected at indicated time points, and viral titers were determined by plaque assay. ***P < 0.001; (one-way ANOVA and Tukey’s test).

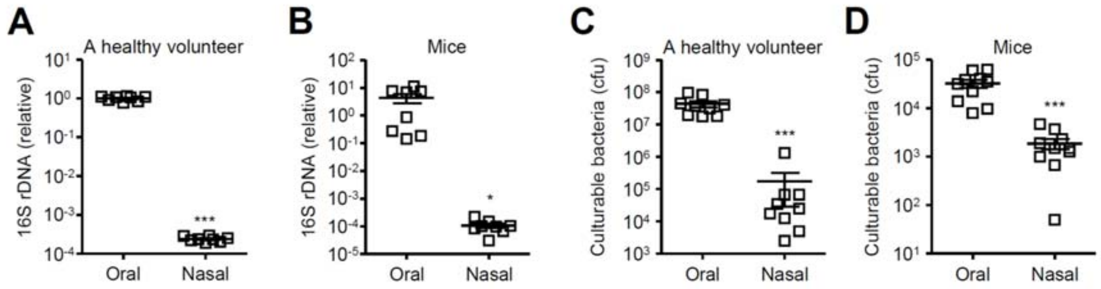


**Figure S3. Relative gene copies of 16S rDNA and bacterial load in in the tongue and nasal wash.** Relative gene copies of 16S rDNA isolated from tongue (A) and nasal wash (B) were quantified by qPCR. Culturable bacterial load in the tongue (C) and nasal wash (D) were measured. *$P < 0.05$ and ***$P < 0.001$; (one-way ANOVA and Tukey’s test).
